# Iron Age and Anglo-Saxon genomes from East England reveal British migration history

**DOI:** 10.1101/022723

**Authors:** Stephan Schiffels, Wolfgang Haak, Pirita Paajanen, Bastien Llamas, Elizabeth Popescu, Louise Lou, Rachel Clarke, Alice Lyons, Richard Mortimer, Duncan Sayer, Chris Tyler-Smith, Alan Cooper, Richard Durbin

## Abstract

British population history has been shaped by a series of immigrations and internal movements, including the early Anglo-Saxon migrations following the breakdown of the Roman administration after 410CE. It remains an open question how these events affected the genetic composition of the current British population. Here, we present whole-genome sequences generated from ten ancient individuals found in archaeological excavations close to Cambridge in the East of England, ranging from 2,300 until 1,200 years before present (Iron Age to Anglo-Saxon period). We use present-day genetic data to characterize the relationship of these ancient individuals to contemporary British and other European populations. By analyzing the distribution of shared rare variants across ancient and modern individuals, we find that today’s British are more similar to the Iron Age individuals than to most of the Anglo-Saxon individuals, and estimate that the contemporary East English population derives 30% of its ancestry from Anglo-Saxon migrations, with a lower fraction in Wales and Scotland. We gain further insight with a new method, rarecoal, which fits a demographic model to the distribution of shared rare variants across a large number of samples, enabling fine scale analysis of subtle genetic differences and yielding explicit estimates of population sizes and split times. Using rarecoal we find that the ancestors of the Anglo-Saxon samples are closest to modern Danish and Dutch populations, while the Iron Age samples share ancestors with multiple Northern European populations including Britain.

---

Within the last 2,000 years alone, the British Isles have received multiple well-documented immigrations. These include military invasions and settlement by the Romans in the first century CE, peoples from the North Sea coast of Europe collectively known as the Anglo-Saxons between ca. 400CE and 650CE (Figure 1a), Scandinavians during the late Saxon “Viking period” 800-1000CE and the Normans in 1066CE^1^. These events, along with prior and subsequent population movements, have led to a complex ancestry of the current British population. Although there is only a slight genetic cline from north to south at a coarse level^2,3^, recent analyses have revealed considerable fine-scale genetic structure in the Northern and Western parts of Great Britain, alongside striking homogeneity in Southern and Eastern England^4^ in the regions where archaeologists identify early Anglo-Saxon artifacts, cemeteries and communities. A variety of estimates of the fraction of Anglo-Saxon genetic ancestry in England have been given^5^-^7^, with the recent fine structure analysis suggesting most likely 10-40%^4^.

**Figure 1:**
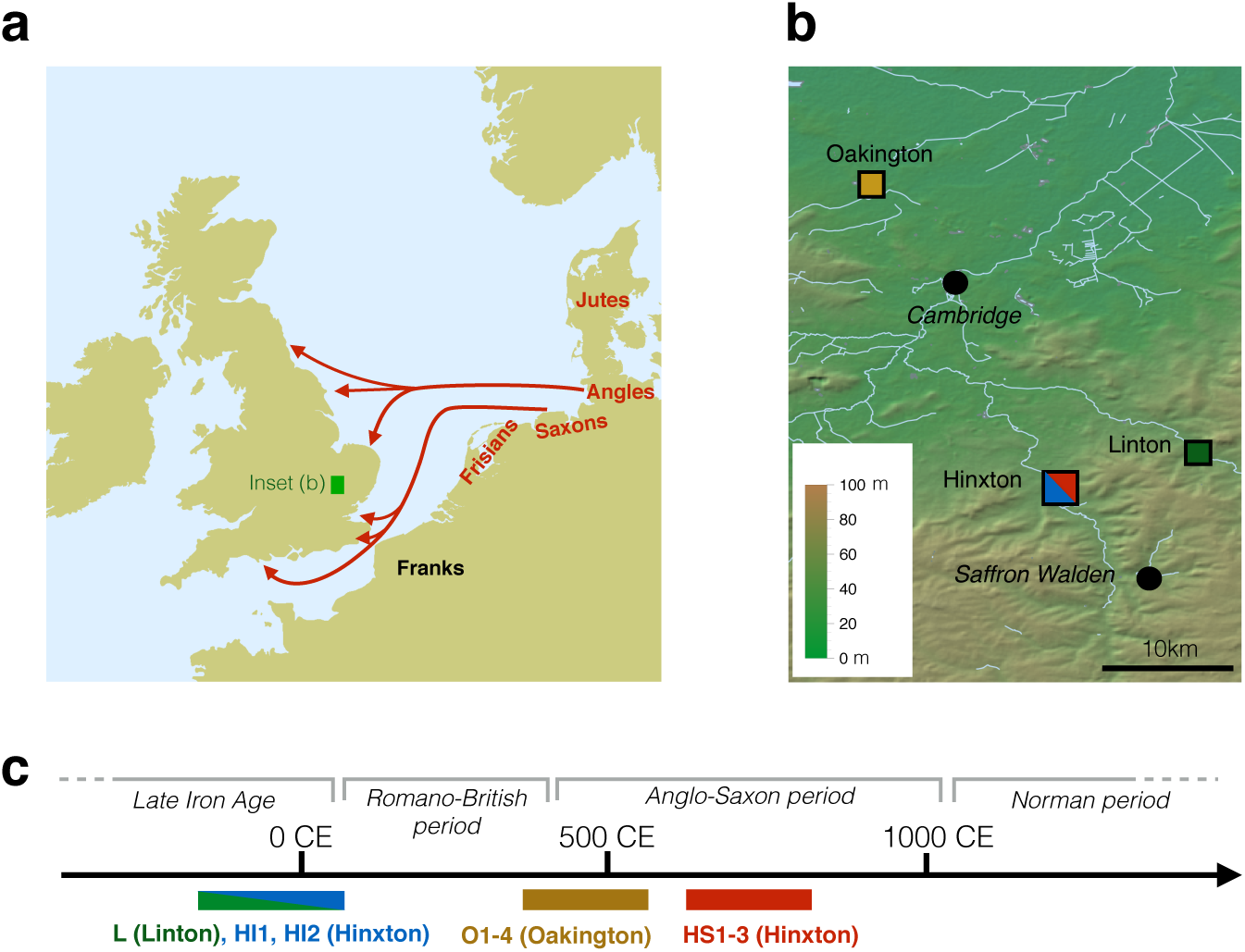
Geographic and temporal context of the samples used in this study. (a) Anglo-Saxon migration routes of people from the continental coast, as reconstructed from historical and archaeological sources ^1,4^. (b) The ancient samples used in this study were excavated at three archaeological sites in East England: Hinxton, Oakington and Linton. The towns Cambridge and Saffron Walden are also shown (black circles). Background green/brown shades indicate altitude. The colors of the four sample match the ones in panel c and Figure 2. (c) The 10 ancient samples belong to 3 age groups. The sample from Linton and two samples from Hinxton are from the late Iron Age, the four Oakington samples from the early Anglo-Saxon period, and three Hinxton samples are from the middle Anglo-Saxon period.

Here we present whole genome sequences of ten ancient samples, excavated in three sites in East England close to Cambridge: Hinxton (5 samples), Oakington (4 samples) and Linton (1 sample) (Figure 1b, Extended Data Figures 1 and 2). All samples were radiocarbon dated (Supplementary Information section 1 and 2), and fall into three time periods: the Linton sample and two Hinxton samples are from the late Iron Age (around 100 BCE), the four samples from Oakington from the early Anglo-Saxon period (5^th^ to 6^th^ century), and three Hinxton samples from the middle Anglo-Saxon period (7^th^ to 9^th^ century) (Figure 1c). The two Iron Age samples from Hinxton are male, all other samples are female, based on Y chromosome coverage. All samples were sequenced to genome wide coverage from 1x to 12x (Table 1). All have contamination rates below 2%, as estimated both from mitochondrial DNA and from nuclear DNA (Extended Data Table 1, Supplementary Information section 5). Mitochondrial and Y chromosome haplogroups of all samples are among the most common haplogroups in present-day Britain (Table 1)^8,9^.

**Table 1:** A summary of all samples in this study. A summary of the samples in this study. The “% endogenous” values give the percentage of sequenced DNA that map to the human reference genome. The “% unique” values give the fraction of mapped reads that are left when excluding duplicates. The “mean autosomal coverage” is the number of reads covering a base, averaged across chromosome 20.

| Name | Origin | Sex | C14 Date | % endogenous | % Unique | MT and Y Haplogroup | mean autosomal coverage |
| --- | --- | --- | --- | --- | --- | --- | --- |
| L | Linton | female | 360 - 50 BCE | 72 % | 54% | H1e | 1.4 |
| HI1 | Hinxton | male | 160 BCE - 26 CE | 16 % | 63% | K1a1b1b,<br>R1b1a2a1a2c | 1.3 |
| HI2 | Hinxton | male | 170 BCE - 80 CE | 83 % | 65% | H1ag1,<br>R1b1a2a1a2c1 | 11.8 |
| O1 | Oakington | female | 420 - 570 CE | 81 % | 50% | U5a2a1 | 3.8 |
| O2 | Oakington | female | 385 - 535 CE | 92 % | 68% | H1g1 | 2.7 |
| O3 | Oakington | female | 395 - 540 CE | 95 % | 64% | T2a1a | 8.2 |
| O4 | Oakington | female | 400 - 545 CE | 67 % | 77% | H1at1 | 6.3 |
| HS1 | Hinxton | female | 666 - 770 CE | 36 % | 91% | H2a2b1 | 4.4 |
| HS2 | Hinxton | female | 631 - 776 CE | 42 % | 74% | K1a4a1a2b | 3.8 |
| HS3 | Hinxton | female | 690 - 881 CE | 16 % | 71% | H2a2a1 | 0.9 |

We generated a principal component plot of the ten ancient samples together with relevant European populations selected from published data^10,11^ (Extended data Figure 3). The ancient samples fall within the range of modern English and Scottish samples, with the Iron Age samples from Hinxton and Linton falling closer to modern English and French samples, while most Anglo-Saxon era samples are closer to modern Scottish and Norwegian samples. Overall, though, population genetic differences between these samples at common alleles are very slight.

While principal component analysis can reveal relatively old population structure, such as generated from long-term isolation-by-distance models^12^, whole genome sequences let us study rare variants to gain insight into more recent population structure. We identified rare variants with allele frequency up to 1% in a reference panel of 433 European individuals from modern Finland, Spain, Italy, Netherlands and Denmark, for which genome-wide sequence data are available ^13^-^15^. We determined for each ancient sample the number of rare variants shared with each reference population (Supplementary Table 1). There are striking differences in the sharing patterns of the samples, illustrated by the ratio of the number of rare alleles shared with Dutch individuals to the number shared with Spanish individuals (Figure 2a). The middle Anglo-Saxon samples from Hinxton (HS1, HS2, HS3) share relatively more rare variants with modern Dutch than the Iron Age samples from Hinxton (HI1, HI2) and Linton (L). The early Anglo-Saxon samples from Oakington are more diverse, with O1 and O2 being closer to the middle Anglo-Saxon samples, O4 exhibiting the same pattern as the Iron Age samples, and O3 showing an intermediate level of allele sharing, suggesting mixed ancestry. The differences between the samples are highest in low frequency alleles and decrease with increasing allele frequency. This is consistent with mutations of lower frequency on average being younger, reflecting more recent distinct ancestry, compared with higher frequency mutations reflecting older shared ancestry.

**Figure 2:**
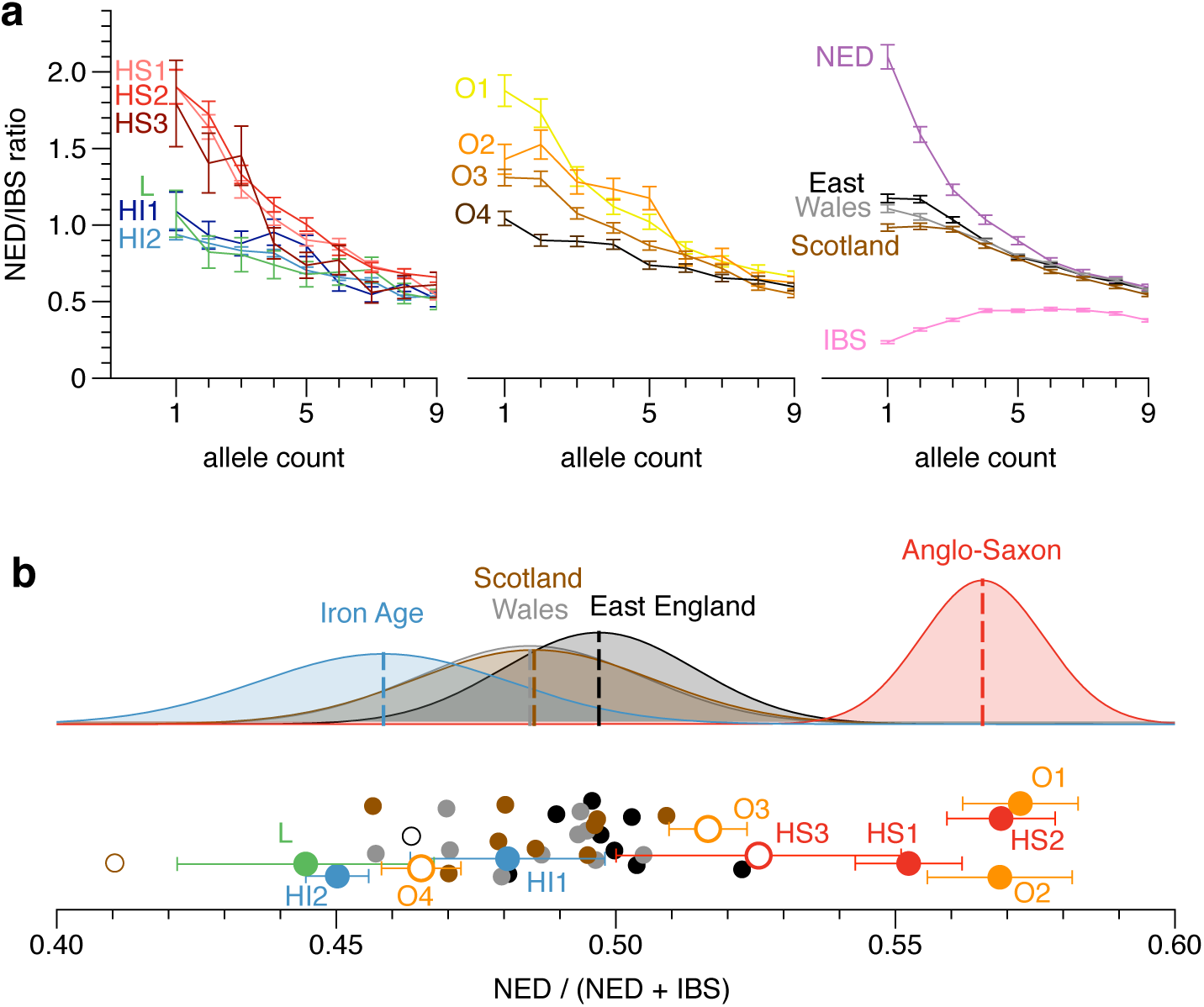
Relative rare allele sharing between ancient and modern samples. (a) The ratio of the numbers of rare alleles shared with modern Dutch and Spanish samples as a function of the allele count in the set of modern samples. Ancient sample codes (left-hand and middle sections) are defined in Table 1. Results from present-day British individuals (right hand panel) are averaged over 10 individuals from each subpopulation. Results from a Dutch and a Spanish individual are shown for comparison. (b) The relative fraction of rare alleles shared with modern Dutch compared to Spanish alleles, integrated up to allele count 5. Iron Age and Anglo-Saxon samples mark the two extremes on this projection, while modern samples are spread between them, indicating mixed levels of Anglo-Saxon ancestry, which is on average higher in East England than elsewhere in Wales and Scotland. Empty symbols indicate that the sample has been excluded from computing the averages indicated above due to being apparently admixed or otherwise anomalous. Samples are shown with a random vertical offset for better clarity. Error bars for the modern samples are omitted here, but of the same order as for the ancient samples.

We also examined using the same method 30 modern samples from the UK10K project ^16^, 10 each with birthplaces in East England, Wales and Scotland. Overall, these samples are closer to the Iron Age samples than to the Anglo-Saxon era samples (Figure 2a). There is a small but significant difference between the three modern British sample groups, with East English samples sharing slightly more alleles with the Dutch, and Scottish samples looking more like the Iron Age samples. To quantify the ancestry fractions, we fit the modern British samples with a mixture model of ancient components, by placing all the samples on a linear axis of relative Dutch allele sharing that integrates data from allele counts one to five (Figure 2b). By this measure the East England samples are consistent with 30% Anglo-Saxon ancestry on average, with a spread from 20% to 40%, and the Welsh and Scottish samples are consistent with 20% Anglo-Saxon ancestry on average, again with a large spread (Supplementary Table 2). An alternative and potentially more direct approach to estimate these fractions is to measure rare allele sharing directly between the modern British and the ancient samples. While being much noisier than the analysis using Dutch and Spanish outgroups, this yields consistent results (Extended Data Figure 4 and Supplementary Table 2). In summary, this analysis suggests that only 20-30% of the ancestry of modern Britons was contributed by Anglo-Saxon immigrants, with the higher number in East England closer to the immigrant source. The difference between the three modern groups is surprisingly small compared to the large differences seen in the ancient samples, although we note that the UK10K sample locations may not fully reflect historical geographical population structure because of recent population mixing.

To get further insight into the history underlying these sharing patterns, we developed a sensitive new method, rarecoal, which fits a demographic model to the joint distribution of rare alleles in a large number of samples (Supplementary Information section 6). The key idea is to model explicitly the uncertainty in the past of the distribution of derived alleles, but approximate the corresponding distribution for non-derived alleles by its expectation (Figure 3a). Because rarecoal explicitly models rare mutations, it estimates separations in mutation clock time rather than genetic drift time, in contrast to methods based on allele frequency changes in common variants^17^. We first tested rarecoal on simulated data and found that it was able to reconstruct split times and branch population sizes with good accuracy (Figure 3b), matching allele sharing almost exactly (Extended Data Figure 5).

**Figure 3:**
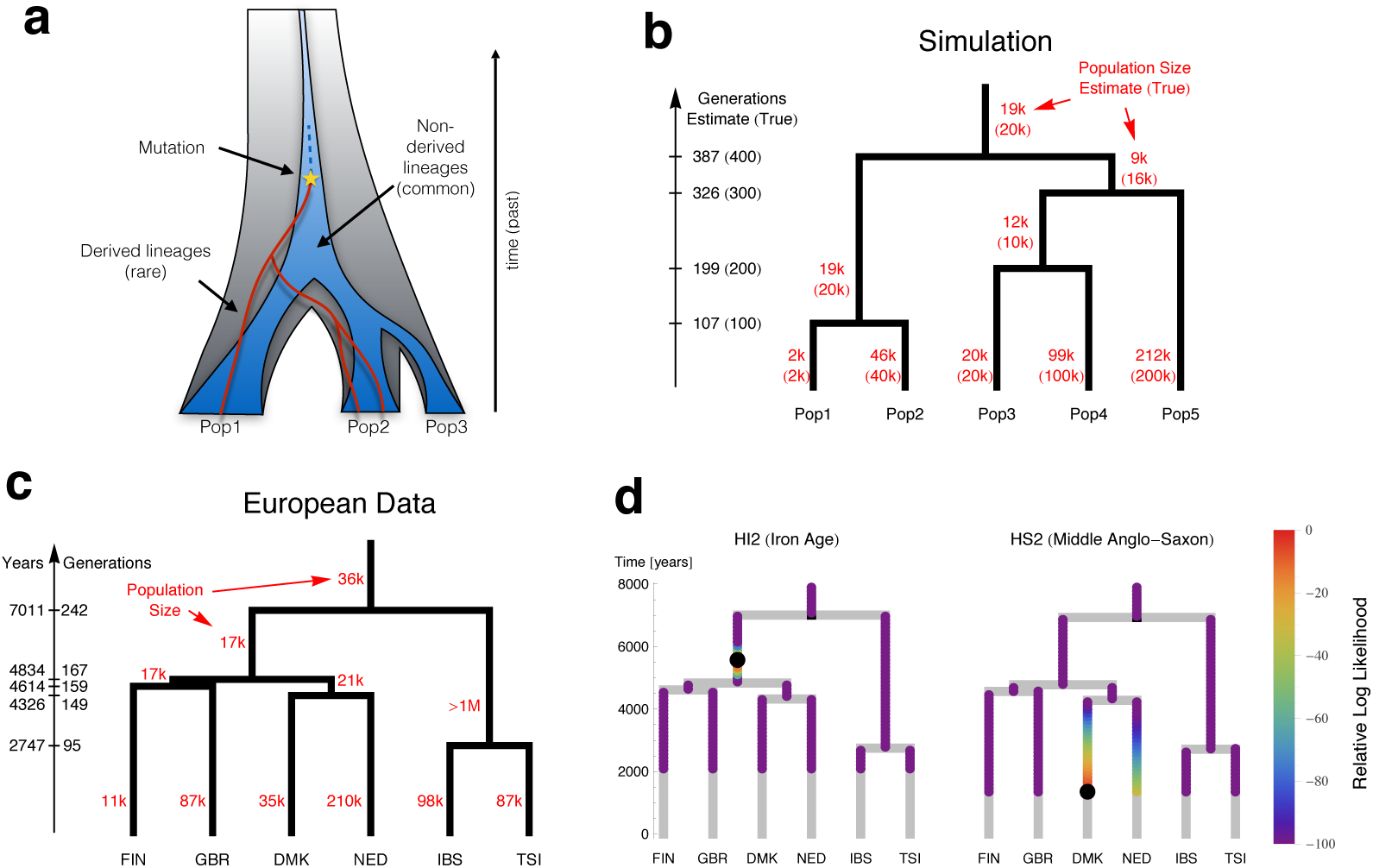
Modeling European history with rarecoal. (a) Rarecoal tracks the probabilities for the lineages of rare alleles (red) within a coalescent framework back in time, and replaces the distribution of non-derived alleles (dark blue) by its average. (b) By optimizing the likelihood of the data under the model, we can estimate population sizes and split times. Tested with simulated data, the estimates closely match the true values (in parentheses). (c) Applied to 524 European individuals, rarecoal estimates split times as indicated on the time axis and population sizes for each branch. (d) Given the European tree, we can map ancient samples onto this tree, illustrated by two samples. We color each point in the tree according the likelihood that the ancestral branch of the ancient sample merges at that point. The maximum likelihood merge point is marked by a black circle.

We next applied rarecoal to 524 samples from six populations in Europe (Figure 3c) to estimate a European demographic tree into which we could place the ancient samples. The first split was between Southern and Northern Europe with a median time around 7,000 years ago, followed by three more separations close in time around 4,500 years ago between Netherlands, Denmark, Finland and Britain. The somewhat surprising clustering of modern British samples with Finns, although close in time to other splits, is probably due to the presence of samples from the Orkney Islands in the British set of samples^13^ (see also below). The timing of the most recent split, between Italy and Spain, around 2,700 years ago, may be a consequence of migration following an earlier separation; the population size of the Italian-Spanish ancestral population was estimated to be extremely large and an upper bound could not be determined, which could be an artifact of ancestral substructure or admixture. The tip branch effective population size is lowest in Finland (∼12,000), consistent with previous observations ^18,19^, and highest in the Netherlands (∼210,000). For the European data, the allele sharing fit is a little less good than for the simulated data (Extended Data Figure 6), presumably due to simplifying model assumptions such as a constant population size in each branch and the absence of migration.

Following this, we placed our ancient samples into the European tree, by evaluating the likelihood for each ancient sample to merge with every possible point on the tree prior to the date of origin of the sample (Figure 3d, Methods and Supplementary Information section 6). There was a marked difference between the Iron Age and the Anglo-Saxon era samples: the Anglo-Saxon era samples mostly merged onto the Dutch and Danish branches, whereas the Iron Age samples preferentially merged at the base of the ancestral branch for all modern Northern European samples (Figure 3d, Extended Data Figure 7). The exception is that the early Anglo-Saxon O4 shows the same signal as the Iron Age samples, consistent with the rare allele sharing analysis (Figure 2). There is some differentiation amongst the Anglo-Saxon era samples, with early Anglo-Saxon sample O1 and O2 having highest likelihood of merging onto the Dutch branch while O3 and the middle Anglo-Saxon HS1, HS2 and HS3 have highest likelihoods of merging onto the Danish branch. The signals from HS3, HI1 and L are more spread due to low coverage, but consistent with the other results. Interestingly, when we placed modern samples onto the tree using the same method, samples from most countries placed on the tip of their respective branch, but GBR samples, collected from Kent, Cornwall and Orkney as part of the Peoples of the British Isles collection^4,20^, placed at varied positions, with some at the top of the northern European subtree near the Iron Age samples (Extended Data Figure 8, Supplementary Information section 6).

The genetic analyses described above add significantly to our picture of Anglo-Saxon migration into Britain. In the cemetery at Oakington we see evidence even in the early Anglo-Saxon period for a genetically mixed but culturally Anglo-Saxon community^21,22^, in contrast to claims for strong segregation between newcomers and indigenous peoples^7^. The genomes of two sequenced individuals are consistent with them being of recent immigrant origin, from different continental source populations, one was genetically similar to native Iron Age samples, and the fourth was an admixed individual, indicating intermarriage. Despite this, their graves were conspicuously similar, with all four individuals buried in flexed position, and with similar grave furnishing. Interestingly the wealthiest grave, with a large cruciform brooch, belonged to the individual of native British ancestry (O4), and the individual without grave goods was one of the two genetically “foreign” ones (O2), an observation consistent with isotope analysis at West Heslerton which suggests that new immigrants were frequently poorer ^23,24^. Given this mixing apparent around 500CE, and that the modern population is no more than 30% of Anglo-Saxon ancestry, it is perhaps surprising that the middle Anglo-Saxon individuals from the more dispersed field cemetery in Hinxton all look genetically consistent with unmixed immigrant ancestry. One possibility is that this reflects continued immigration until at least the Middle Saxon period. The unmixed Hinxton group, versus the mixing of the Oakington population, shows that early medieval migration took a variety of forms and that these migrants integrated with the incumbent population in different ways. Full genome sequences, and new methods such as rarecoal, now allow us to use slight distinctions in genetic ancestry to study such recent events. Further ancient genomes, and methodological improvements to incorporate explicit migration and mixing, will enable us to resolve them in more detail.

## Acknowledgements

We thank everyone who contributed to the archaeological excavations, the sequencing team at the Wellcome Trust Sanger Institute, and David Reich’s laboratory for contributing to the characterization of the libraries. We thank Luka Papac for wet lab support at the Australian Centre for Ancient DNA. The Oakington excavations were funded by the Oakington Parish Council, the Institute for Field Research (IFR) and the University of Central Lancashire (UCLan). This work was funded by Australian Research Council grant DP130102158, by the University of Adelaide’s Environment Institute, and by Wellcome Trust Grant 098051.

## Author Contributions

SS, AC, CTS and RD designed and oversaw the study. EP, RC, AL provided samples from Linton and Hinxton, DS provided samples from Oakington. WH prepared samples and extracted DNA, WH and BL generated sequencing libraries. SS, PP and RD developed methods. SS, PP and RD analyzed data. EP, RC, AL, LL, RM and DS provided archaeological context. SS, RD and DS wrote the paper and all contributed comments.

## Author information

The raw sequence data of the 10 samples presented in this paper are deposited at the European Nucleotide Archive (http://www.ebi.ac.uk/ena). The study IDs are ERP003900 (Hinxton samples) and ERP006581 (Oakington and Linton samples).

The authors have no competing interests.

## Methods

Custom software mentioned here is publically available on <u>www.github.com/stschiff/sequenceTools</u> if not stated otherwise.

### Library screening and sequencing

DNA was extracted in clean room facilities in Adelaide using an in-solution silica-based protocol ^25^. Libraries were generated from the Hinxton individuals (n=6) with ^26^ and without enzymatic damage repair (Supplementary Table 3), whereas partial damage repair ^27^ was performed for the Linton (n=3) and Oakington (n=14) samples. All 29 libraries were prepared with truncated barcoded Illumina adapters and amplified with full-length indexed adapters for sequencing^28^. The libraries were screened for complexity and endogenous DNA on an Illumina MiSeq platform in Harvard in collaboration with David Reich. Ten libraries were selected based on high endogenous fractions and high complexity, and sent to the Wellcome Trust Sanger Institute for whole genome sequencing. We sequenced the 10 libraries on a total of 52 lanes on an Illumina HiSeq 2500 platform with a 75bp paired end protocol.

### Raw sequencing data processing

Because typical ancient DNA inserts are about 50bp long, we expected significant overlaps between the read pairs. We aligned reads in each pair, discarded non overlapping pairs, and merged paired reads that overlapped by at least 30bp before removing barcodes and adaptors using our custom program “filterTrimFastq". The resulting collection of merged reads was aligned to the Human Reference build 37 using bwa ^29^ for each sample. Alignments were sorted and duplicates removed with our custom script “samMarkDup.py". We assessed low levels of ancient DNA damage, present despite damage repair, in the aligned reads using mapDamage2 ^30^ and rescaled base qualities using mapDamage2.

### Mitochondrial and Y chromosome analysis

We called mtDNA and Y chromosome consensus sequences using samtools. Haplogroups were handcurated using public databases (Supplementary Information section 5, Supplementary Table 4).

### Contamination Estimates

We estimated possible modern DNA contamination in all ancient samples using two methods. First, we tested for evidence for contaminant mitochondrial DNA^31^. We looked for sites in the mitochondrial genome, at which the ancient sample carried a consensus allele that was rare in the 1000 Genomes reference panel. We then looked whether there were reads at these sites that carried the majority allele from 1000 Genomes (Supplementary Information section 5). Second, we used the program “verifyBamId” ^32^ to carry out a similar test in the nuclear genome, again using the 1000 Genomes reference panel. Contamination estimates are summarized in Extended Data Table 1.

### Principal component analysis

We downloaded the Human Origins Data set ^10,11^ and called genotypes at all sites in this data set for all ancient samples using a similar calling method as described in ^11^: Of all high quality reads covering a site, we picked the allele that is supported by the majority of reads, requiring at least two reads supporting the majority allele, otherwise we call a missing genotype. If multiple alleles had the same number of supporting reads, we picked one at random. Principal component analysis was performed using the smartpca program from EIGENSOFT ^33^, by using only the modern samples for defining the principal components and projecting the 10 ancient samples onto these components.

### Rare allele sharing analysis

We generated a reference panel consisting of 433 individuals from Finland (n=99), Spain (n=107), Italy (n=107), Netherlands (n=100) and Denmark (n=20). The Finnish, Spanish and Italian samples are from the 1000 Genomes Project (phase 3) ^13^, the Dutch samples from the GoNL project ^14^ and the Danish samples from the GenomeDK project^15^. For the Dutch and Danish samples, only allele frequency data was available. In case of the Dutch data set, we downsampled the full data set to obtain the equivalent of 100 samples. All other reference sample variant calls were used as provided by the 1000 Genomes Project. In addition, we filtered based on a mappability mask ^34,35^ that is available from <u>www.github.com/stschiff/msmc</u>. We selected all variants up to allele count 9 in this reference set and tested for each ancient individual and each of those sites whether the ancient individual carried the rare allele. We called a rare variant (always assumed heterozygous) in the ancient sample if at least two reads supported the rare allele from the reference set. While this calling method will inevitably miss variants in low coverage individuals, the relative numbers of shared alleles with different populations is unbiased.

We accumulated the total number of alleles shared between each ancient sample and each modern reference population, and stratified by allele count in the reference population, up to allele count 9 (Supplementary Table 1). We found that sharing with the Dutch and the Spanish population showed the largest variability across the ancient samples. So for the plot in Figure 2a, we divided the sharing count with the Dutch population by the sharing count of the Spanish population for each allele frequency. To plot curves from the Dutch and the Spanish population itself, we sampled haploid individuals from each population, by sampling with replacement at every variant site in the reference set. This was necessary because for the Dutch samples no genotype information was publically available, only allele frequency data.

For the 30 UK10K samples shown in Figure 2a and b, we started from the read alignment for each individual and called rare variants with respect to the 433 reference individuals in exactly the same way as we did for the ancient samples. For Figure 2a, the allele sharing counts were then accumulated across the 10 individuals in each group.

Error bars for each allele sharing count are based on the square root of each count. For Figure 2b we added the allele sharing counts between each ancient sample and each reference population up to allele count 5, and computed the ratio *NED*/(*NED* + *IBS*), where *NED* is the sharing count with Dutch, and *IBS* the sharing count with Spanish. For the mean and variances shown in Figure 2b, we excluded outliers as indicated in the caption of the Figure. The fraction of Anglo-Saxon derived ancestry is computed for each modern UK10K sample as the relative distance of its relative sharing ratio from the Iron Age mean value compared to the Saxon era mean value, as shown in Figure 2b, with 0% corresponding to the Iron Age mean, and 100% corresponding to the Anglo-Saxon era mean.

A similar calculation was done for Extended Data Figure 2, where we took the entire TwinsUK data set from UK10K (with genotype calls provided by UK10K), consisting of 1854 individuals from across the UK, as a reference panel and computed allele sharing of each ancient sample with subpopulations from Wales, East England and Scotland, using all variants up to allele count 37 (1%) in the full data set. In this case, because we had to normalize out coverage differences between the ancient samples, we divided the sharing counts for each ancient sample by the number of shared variants with TwinsUK with allele counts 37 through 370 (1%-10%). We then computed for each TwinsUK sample the mean normalized sharing count with the Iron Age group (H1, H2 and L) and with the Anglo-Saxon era group (HS1, HS2, O1 and O2). We did the same calculation for each ancient individual, by first removing that individual from the two groups above and comparing to the rest of each group, with the same outliers removed as for Figure 2b.

### The Rarecoal method

Rarecoal is a new framework to calculate the joint allele frequency spectrum across multiple populations using rare alleles. Given a certain distribution of rare derived alleles across subpopulations (here up to allele count 4), and a given number of non-derived alleles, which can be arbitrarily large, we want to calculate the total probability of that configuration under a demographic model. The model consists of a population tree with constant population sizes in each branch of the tree, and split times. To give rise to the data observed in the present, the lineages of the derived alleles must coalesce among each other before they coalesce to any non-derived lineage. We introduce a state space that contains all possible configurations of derived lineages across populations and propagate a probability distribution over this space back in time.

We initialize this probability distribution with the given configuration in the present. Time is continuous and discretized into intervals that correspond to one generation in the beginning and cross over to exponentially increasing intervals further in the past. As we go back into the past, we use the structured coalescent to update the state probabilities using the population sizes and split times of the model, until all derived lineages have with very high relative probability coalesced into one. For these updates we need to know the number of non-derived lineages in each population at each time, for which we introduced a major simplification: Instead of modeling the full probability distribution over different numbers of non-derived lineages in each population, we used the expected number of non-derived lineages in each population as a deterministic variable through time. This can be easily updated for each time interval, alongside the state space updates. This way, we keep the probability space small, which makes this method scalable to an arbitrary number of samples, as long as the number of derived alleles is small.

Ultimately, we compute the expected branch length, *d*(***m***; Θ), given the starting configuration *C* and the model Θ, on which the mutation must have happened that gave rise to the pattern that we see in the present. The likelihood *L*(***m***|Θ) of the observed configuration, ***m***, given the model *M*, is then *L*(***m***|Θ) = *d*(***m***; Θ) θ/ *2*, where *θ* = 4*Nμ* is the scaled mutation rate. Given a joint allele frequency histogram with counts *n*(***m***_*i*_) for each rare allele configuration ***m***_*i*_, the total log likelihood of the data is then

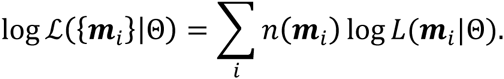

The configurations ***m***_*i*_ consist of joint allele counts across multiple populations up to some frequency, which in our case is allele count 4. See Supplementary Information section 6 for details and derivations.

The outputs from rarecoal are in scaled time. To convert to real time (years) and real population sizes, we used a per-generation mutation rate of 1.25×10^−8^ and a generation time of 29 years.

### Rarecoal analysis

We implemented rarecoal in a program (<u>www.github.com/stschiff/rarecoal.hs</u>) that can learn the parameters of a given population tree topology from the data using numerical maximization of the likelihood and subsequent Markov Chain Monte Carlo to get posterior distributions for each split time and branch population size. We did not implement an automated way to learn the tree topology itself, but use a step by step protocol to learn the best topology fitting the data, adding one population at a time (Supplementary Information section 6).

We tested the method on simulated data using the SCRM simulator ^36^ with the model shown in Figure 3b, with 1000 haploid samples distributed evenly across the 5 populations and realistic recombination and mutation parameters. We then learned the model from the European data set as shown in Figure 3c using a similar protocol.

For mapping ancient samples on the tree we used the same calling method as in the rare allele sharing analysis. We then added the ancient individual as a separate seventh population to the European tree and evaluated the likelihood for this external branch to merge anywhere on the tree. We restricted the fitting to alleles that were shared with the ancient sample. We also made sure that the age of the ancient sample was correctly modeled into the joint 7-population tree, but “freezing” the state probabilities from the present up to the point where the ancient sample lived.

For testing the tree-coloring method, we used single individuals from within the reference set and used them as separate sample to be mapped onto the European tree. (Supplementary Information section 6).

---

**Extended Data Table 1:**
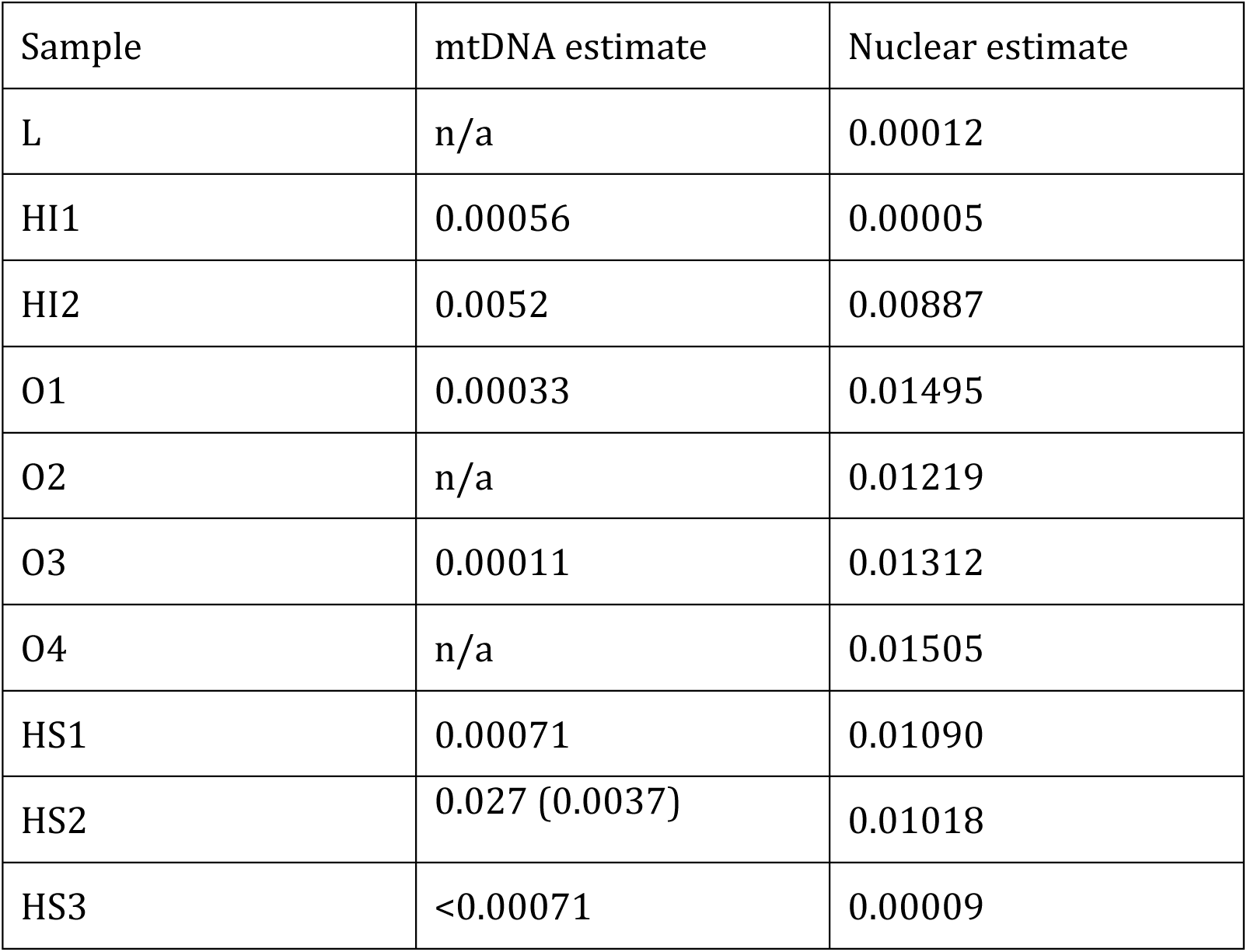
Contamination estimates. DNA contamination estimates based on mitochondrial and nuclear DNA. Numbers are contamination fractions on a 0-1 scale. For O2, O4 and L, no mtDNA estimate could be generated because there were no informative sites. The relatively high contamination estimate of HS2 is due to a single site in the hypervariable region, which could reflect natural heteroplasmy. The estimate without that site is given in parentheses for that individual (Supplementary Information section 5).

**Extended Data Figure 1:**
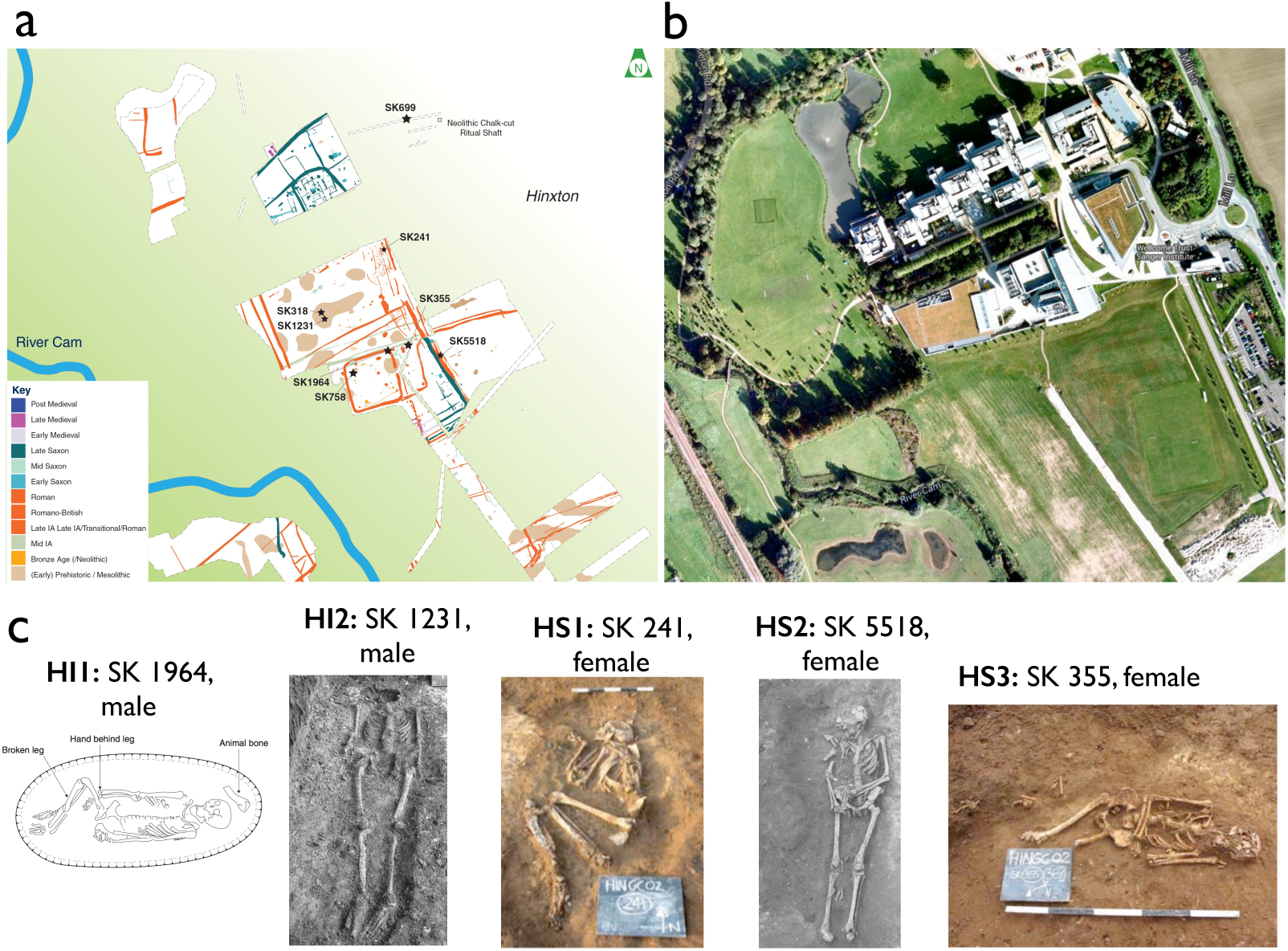
Hinxton Site. (a) A plan of the Hinxton archaeological site, with the locations of the skeletal remains. (b) A satellite image of the same area, where today the Wellcome Trust Genome Campus is located. (c) Pictures/Drawing of the 5 samples used in this study.

**Extended Data Figure 2:**
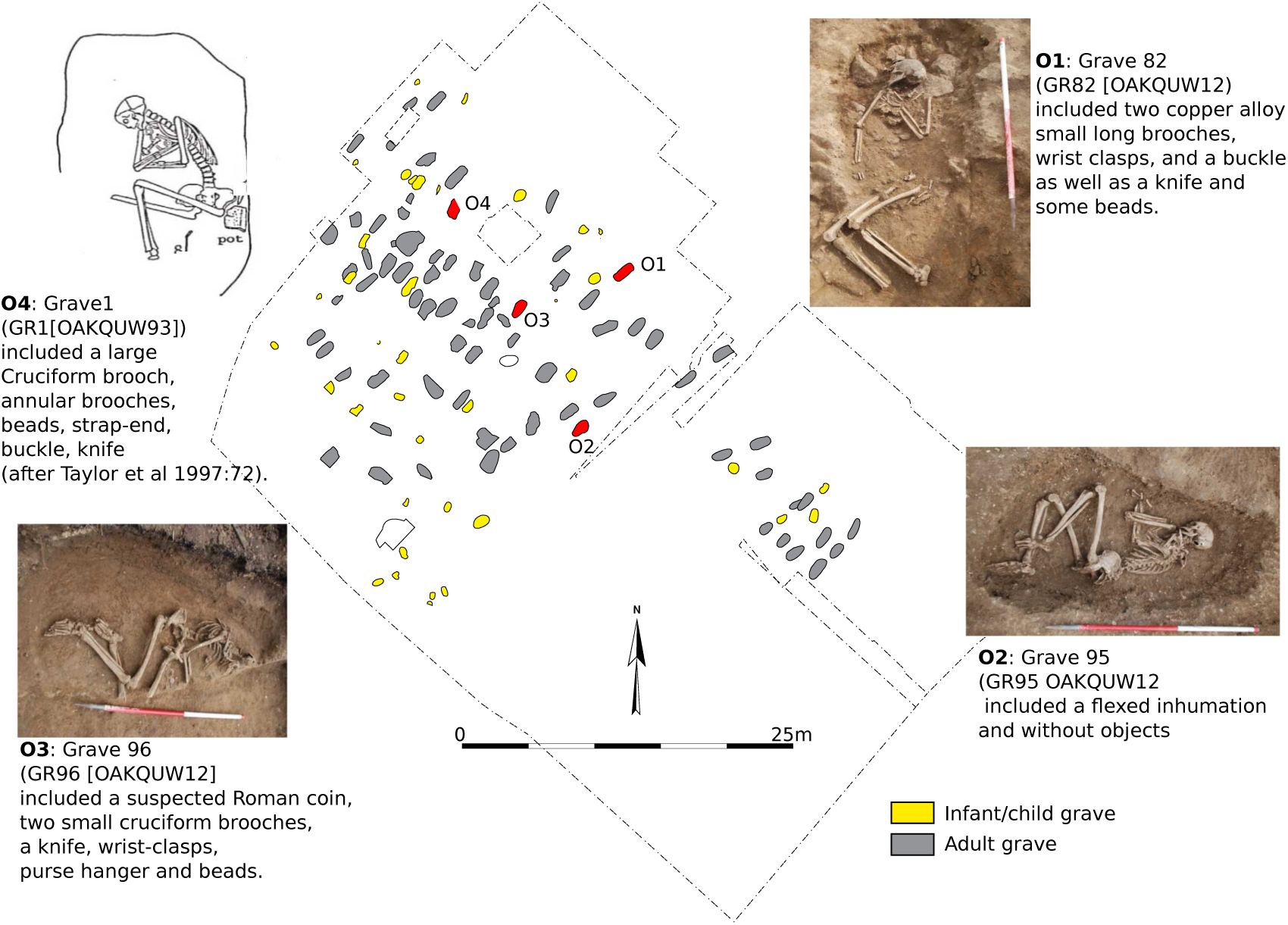
Oakington Site. A schematic of the early Anglo-Saxon cemetery in Oakington, with graves colored in grey (adult individuals), yellow (infant individuals) and red (the adult individuals used in this study).

**Extended Data Figure 3:**
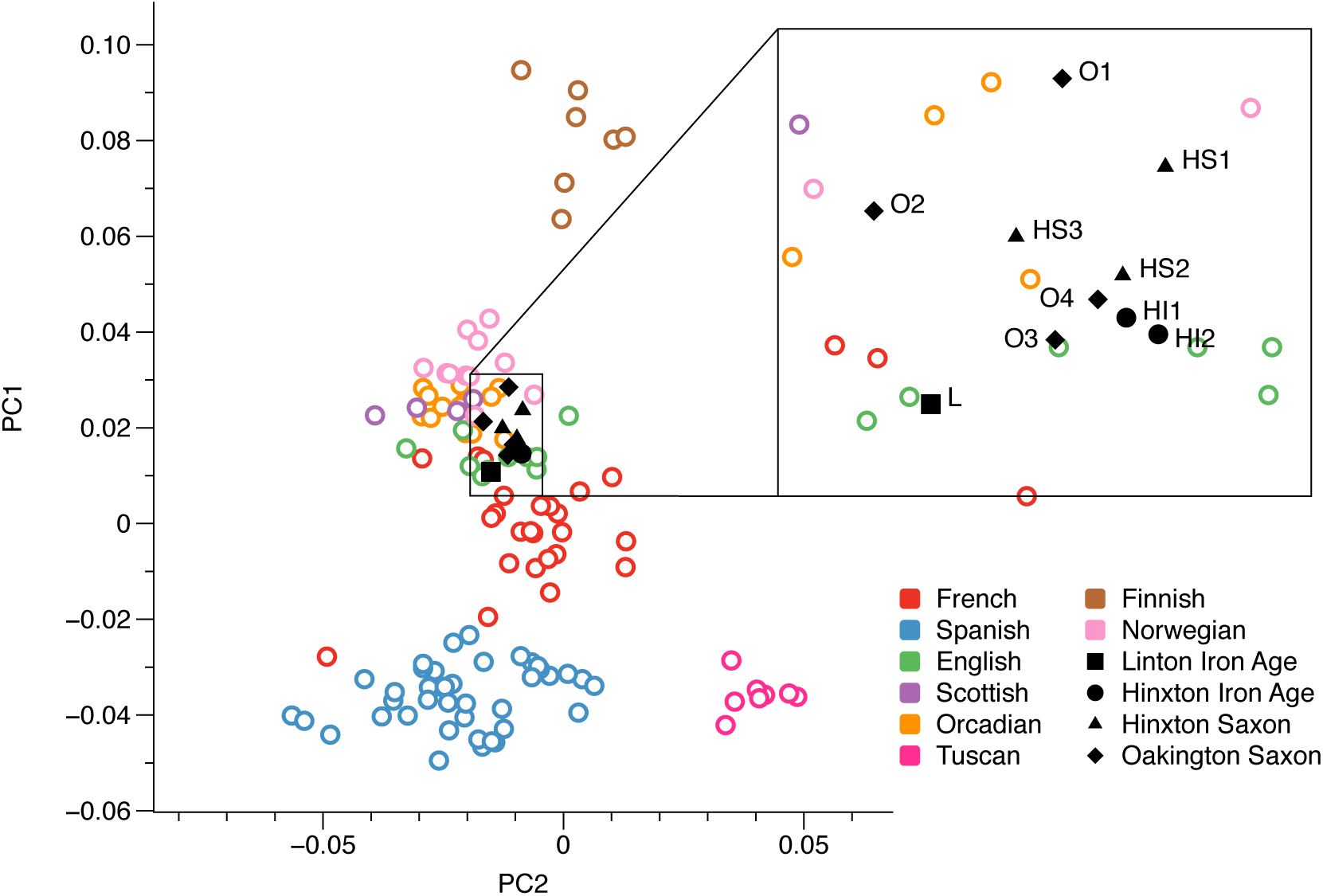
Principal component analysis. The first two principal components obtained by analyzing European samples from the Human Origins Data set ^10,11^ and projecting the ancient samples onto these components.

**Extended Data Figure 4:**
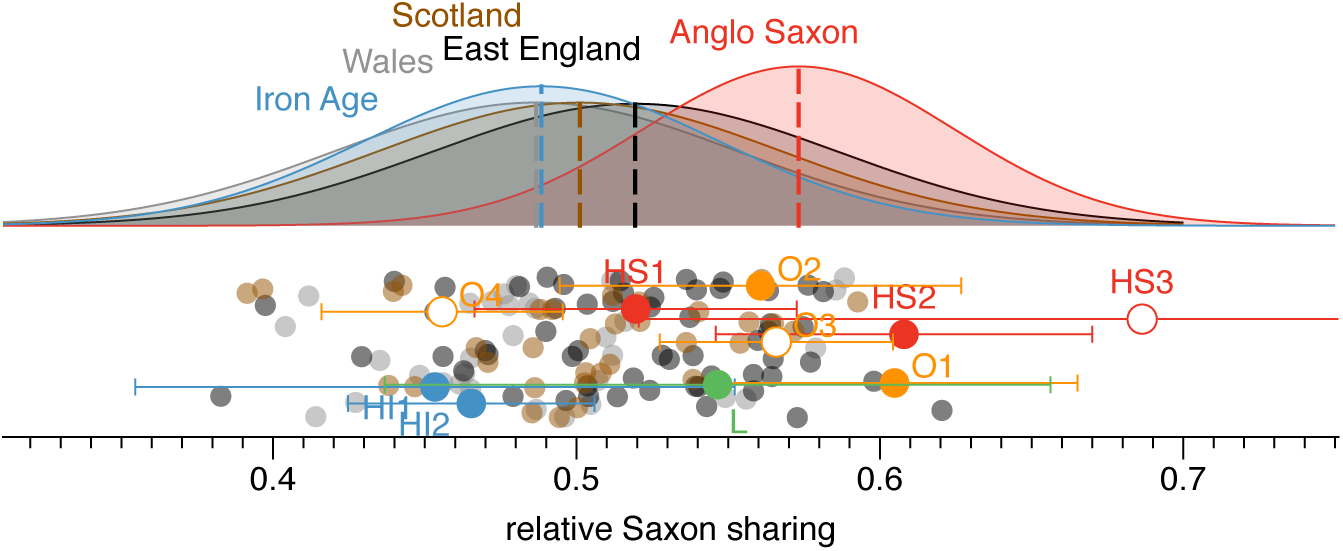
Rare allele sharing of modern British samples with ancient samples. Similarly to Figure 2b, we can obtain rare allele sharing counts between each modern sample and the two age groups of ancient samples, Iron Age and Saxon era. Consistent with Figure 2b, this gives a higher relative Saxon sharing on average of East English samples compared to samples from Wales and Scotland. Because we directly compare each modern sample to a small number of ancient samples, these results are much noisier than the analysis using large Dutch and Spanish outgroup populations as in Figure 2b.

**Extended Data Figure 5:**
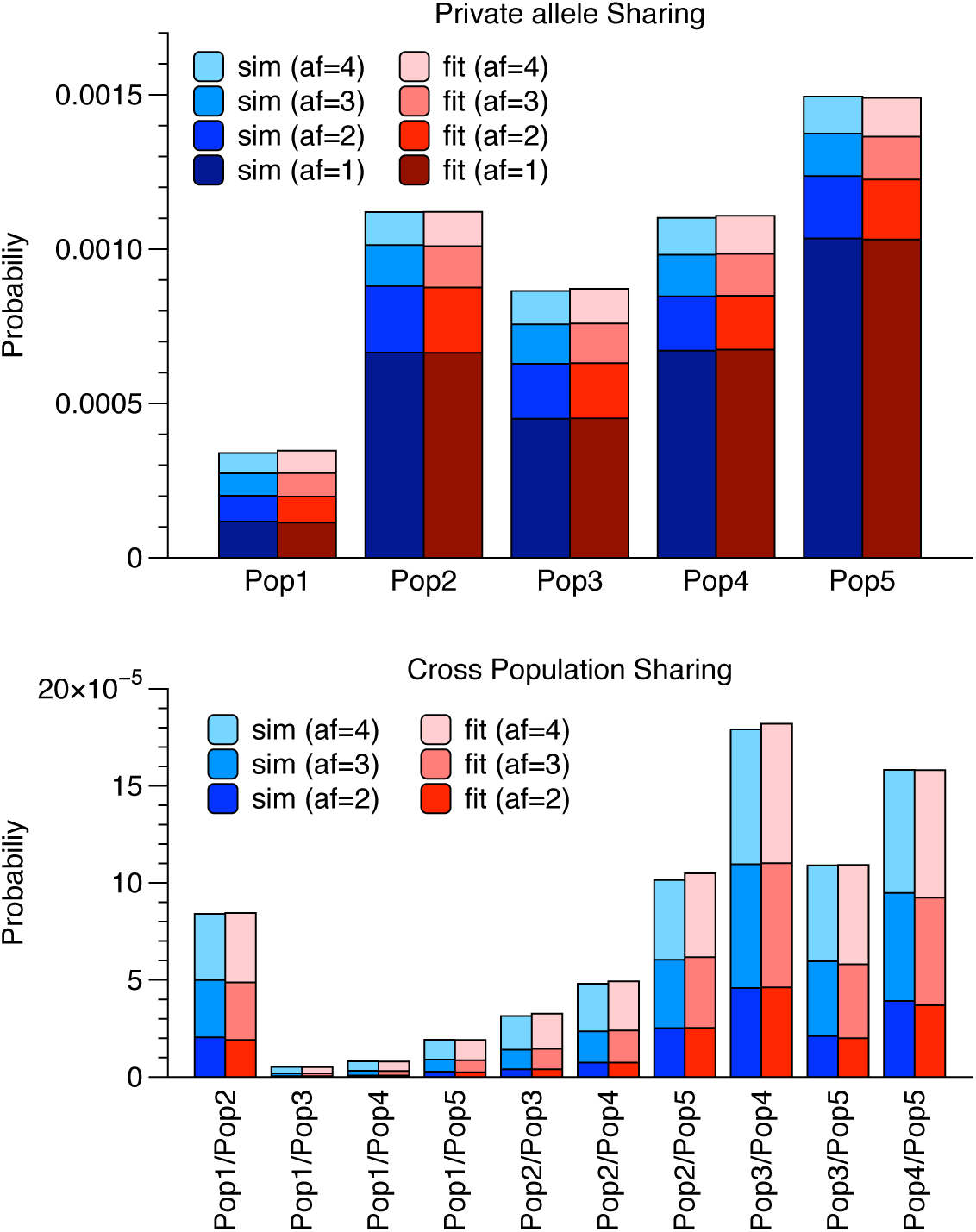
Rarecoal fits of simulated data. We compare the theoretical distribution of rare variants predicted by the model estimated in Figure 3b (red) with the true distribution of variants (blue), yielding a good fit of the model given the data. The top panel shows variants private to one population, the lower panel shows variants shared across populations.

**Extended Data Figure 6:**
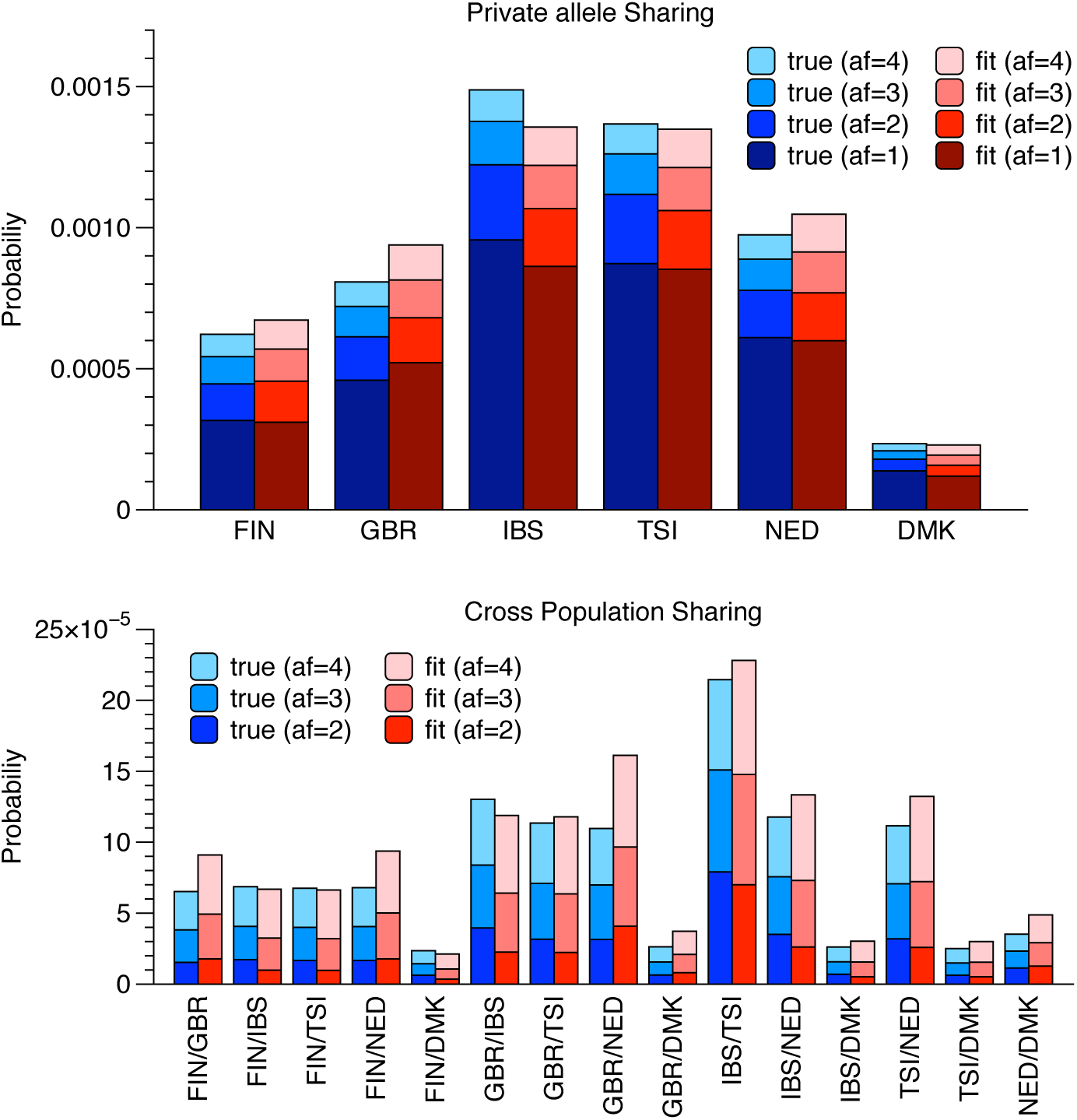
Rarecoal fits of European data. Similar to Extended Data Figure 5, we obtain fits between the model obtained on the European samples (Figure 3c) with the true distribution of rare variants. The fit is reasonable, with some systematic differences owing to simplifying assumptions such as constant population sizes and the absence of migration.

**Extended Data Figure 7:**
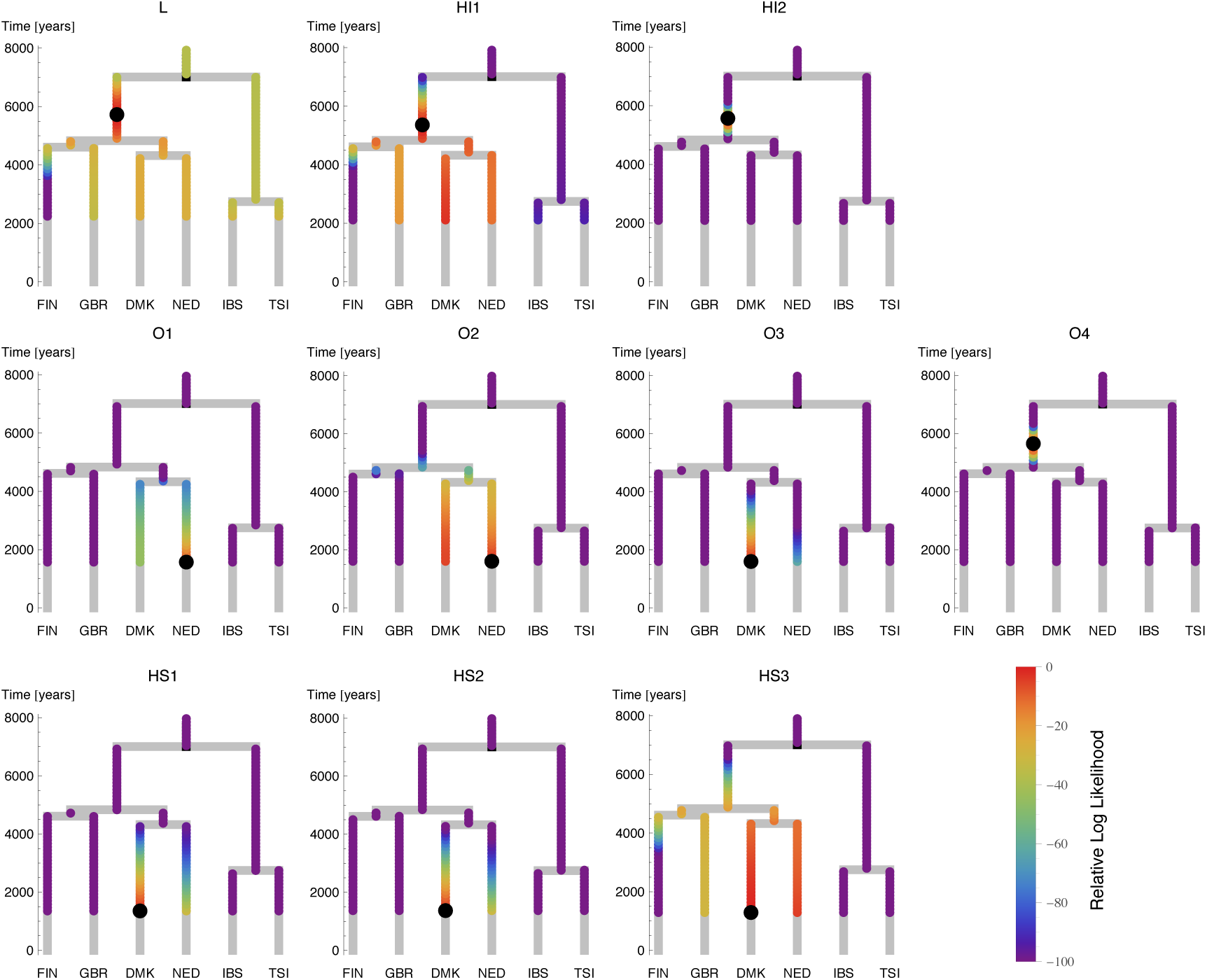
Rarecoal tree painting with ancient samples. The likelihood surface along the tree for each ancient sample, similarly to Figure 3d. Maximum likelihood merge points for each sample are indicated by black circles. For the low coverage samples L, HI1 and HS3 the likelihood is less well localized but consistent with the other samples. We believe the early Anglo-Saxon sample O3 is of mixed immigrant/native ancestry (see Figure 2 and text), but this is not seen here because the rarecoal model assumes a single point to merge onto a tree and thus does not allow for mixed ancestry.

**Extended Data Figure 8:**
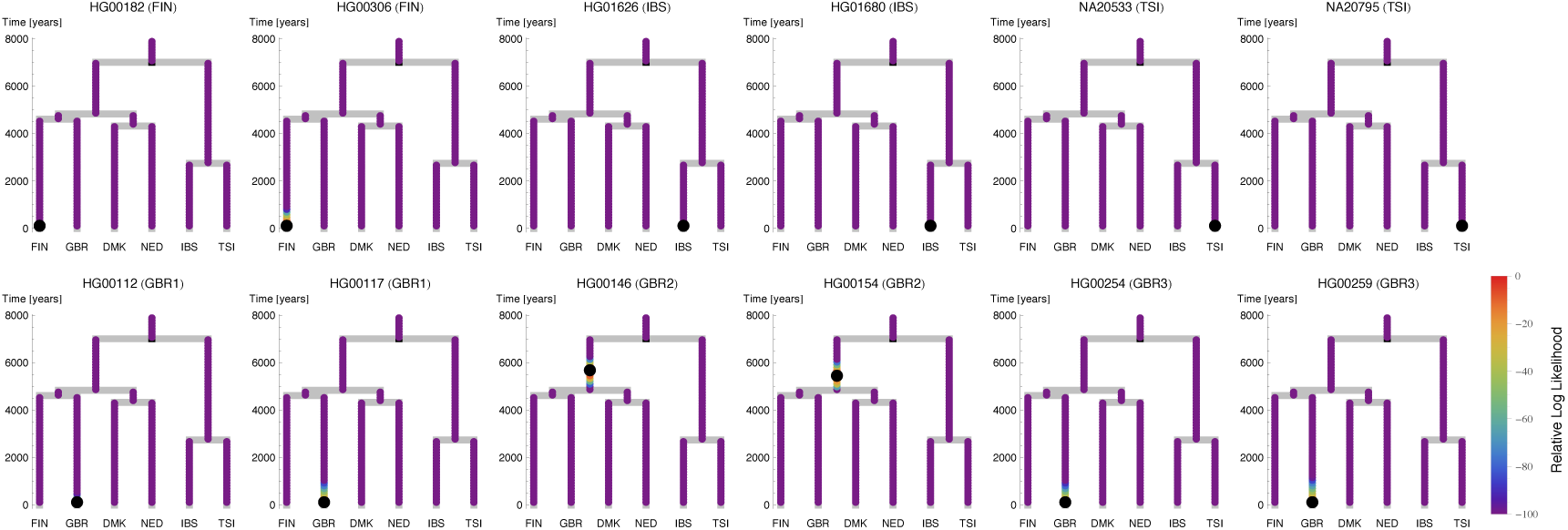
Rarecoal tree painting with modern samples. The likelihood surface along the tree (see Extended Data Figure 7) for several modern samples from the 1000 Genomes project. Most samples map correctly onto the tip of their respective branches, with some GBR samples mapping to the Northern European ancestral branch, due to substructure in the GBR sample set. The black dot indicates the maximum likelihood merge point onto the tree.

